# Urotensin II-Related Peptides, Urp1 and Urp2, Control Zebrafish Spine Morphology

**DOI:** 10.1101/2022.08.13.503856

**Authors:** Elizabeth A. Bearce, Zoe H. Irons, Johnathan R. O’Hara-Smith, Colin J. Kuhns, Sophie I. Fisher, William E. Crow, Daniel T. Grimes

## Abstract

The spine provides structure and support to the body, yet how it develops its characteristic morphology as the organism grows is little understood. This is underscored by the commonality of conditions in which the spine curves abnormally such as scoliosis, kyphosis and lordosis. Understanding the origin of such spinal curves has been challenging in part due to the lack of appropriate animal models. Recently, zebrafish have emerged as promising tools with which to understand the origin of spinal curves. Using zebrafish, we demonstrate that the Urotensin II-related peptides (URPs), Urp1 and Urp2, are essential for maintaining spine morphology. Urp1 and Urp2 are 10-amino acid cyclic peptides expressed by neurons lining the central canal of the spinal cord. Upon combined genetic loss of Urp1 and Urp2, adolescent-onset planar curves manifested in the caudal region of the spine, akin to a lordosis-like condition. Highly similar curves were caused by mutation of Uts2r3, an URP receptor. Quantitative comparisons revealed that Urotensin-associated curves were distinct from other zebrafish spinal curve mutants that more closely reflected idiopathic scoliosis or kyphosis. Last, we found that the Reissner fiber, a proteinaceous thread that sits in the central canal and has been implicated in the control of spine morphology, breaks down prior to curve formation in an idiopathic scoliosis model but was unperturbed by loss of Uts2r3. This suggests a Reissner fiber-independent mechanism of curvature in Urotensin-deficient mutants. Overall, our results show that Urp1 and Urp2 control zebrafish spine morphology and establish new animal models of lordosis-like curves.

## INTRODUCTION

Understanding how the shape of organisms is acquired is a central goal of developmental biology. The chordate body axis forms during embryonic development, when it is based around the rod-like notochord (Stemple, 2005). Later, the vertebrate axis comprises a column of repeating vertebrae which grows during juvenile and adolescent phases and is then maintained during adult life for up to several decades in some species (Bagnat and Gray, 2020). While a great deal has been learned about how the body axis emerges during embryogenesis, less is known about how spine morphology is maintained during growth.

A breakdown of spine morphology occurs in scoliosis, lordosis and kyphosis. Scoliosis is medically defined as lateral curvatures of the spine greater than 10 degrees (Cheng et al., 2015; Mesiti, 2021; Wise et al., 2008) and can be caused by congenital defects of vertebral patterning or as a secondary consequence of neuromuscular disease (Pourquié, 2011; Wishart and Kivlehan, 2021). However, most cases of scoliosis are idiopathic in nature, with no known etiology: approximately 3% of children are afflicted by idiopathic scoliosis, which most often onsets during adolescence (Cheng et al., 2015; Labrom et al., 2021). By contrast, kyphosis and lordosis occur when there is excessive curvature of the thoracic and lumbar regions of the vertebral column, respectively, resulting in a hunched upper back (kyphosis) or a concave lower back (lordosis) (Ogura et al., 2021) without vertebral structural defects. Since these categories of curves can co-occur, there are likely to be overlapping as well as distinct causes.

A challenge to understanding the origin of spinal curvature has been the dearth of suitable animal models recapitulating disease states. Recently, teleost fishes, in particular zebrafish (*Danio rerio*), have emerged as prominent animal models of spinal deformity (Bagnat and Gray, 2021; Bearce and Grimes, 2021; Boswell and Ciruna, 2017; Gorman and Breden, 2009; Roy, 2021). Using zebrafish, it was found that motile cilia-generated cerebrospinal fluid (CSF) flow is essential for maintaining body and spine morphology (Grimes et al., 2016). Mutants with defective motile cilia failed to undergo axial straightening during embryogenesis and so developed a misshapen early embryonic body axis called “curly tail down” (CTD; Brand et al., 1996). If rescued during this early stage, mutants went on to develop three-dimensional spinal curves that recapitulated some features of idiopathic scoliosis, including adolescent-stage onset and absence of vertebral structural defects (Grimes et al., 2016; Marie-Hardy et al., 2021; Wang et al., 2022). Precisely how motile cilia and CSF flow maintain spine morphology during growth is not understood, but it is known that during early larval stages cilia motility is essential for the assembly of the Reissner fiber (RF), an extracellular thread-like structure composed predominantly of the large glycoprotein SCOspondin (encoded by *sspo*) which sits in the CSF in brain ventricles and the central canal (Cantaut-Belarif et al., 2018; Rodriguez et al., 1998). Zebrafish *sspo* mutants exhibited CTD as embryos while hypomorphic mutants which can survive beyond embryonic stages also manifested spinal curves (Cantaut-Belarif et al., 2018; Lu et al., 2020; Rose et al., 2020; Troutwine et al., 2020).

The Urotensin II-related peptides, Urp1 and Urp2, may also function downstream of motile cilia in the central canal. Urp1 and Urp2 are 10-amino acid cyclic peptides previously linked to heart disease and mental illness (Sugo et al., 2003; Konno et al., 2013; Nobata et al., 2011; Parmentier et al., 2011; Quan et al., 2021; Tostivint et al., 2006; Vaudry et al., 2010). In zebrafish, Urp1 and Urp2 are expressed in CSF-contacting neurons (CSF-cNs), flow sensory neurons in the central canal, and their expression is increased by motile cilia function and the RF (Cantaut-Belarif et al., 2020; Lu et al., 2020; Quan et al., 2015; Zhang et al., 2018). Morpholino knockdown of Urp1/Urp2 results in embryonic CTD phenotypes while addition of Urp1/ Urp2 peptides can rescue the CTD of cilia motility- and RF-deficient mutants (Lu et al., 2020; Zhang et al., 2018). This suggested that Urp1 and Urp2 act downstream of cilia motility to promote early axial straightening (Grimes, 2019; Lu et al., 2020; Zhang et al., 2018).

Here, we set out to address whether Urp1 and Urp2 function beyond embryogenesis in maintaining body and spine morphology during growth. By generating zebrafish mutants lacking Urp1 and Urp2 peptides, we found that they are essential, in a mostly redundant fashion, for adult spine morphology. Loss of Urp1 and Urp2 led to the onset of spinal curves during adolescent stages and, by adulthood, resulted in planar curves in the caudal region of the spine that occurred without vertebral structural malformations. A similar phenotype was present upon mutation of the Urotensin receptor gene, *uts2r3*, suggesting that Urp1 and Urp2 signal via Uts2r3 to maintain spine morphology. Urotensin-associated curves were quantitatively distinct from the curves displayed by *cfap298* mutants, which lack cilia motility, and *pkd2l1* mutants in which a CSF-cN-localized ion channel is mutated. Moreover, we found that RF breakdown preceded curve formation in *cfap298* mutants but that RF structure was maintained before and after curves appeared in Urotensin mutants. Overall, this demonstrates that Urp1 and Urp2 peptides control the morphology of the zebrafish spine. We suggest that Urotensin-deficient mutants model a lordosis-like condition and will be important tools for deciphering how the spine is maintained and how this process goes wrong in disease.

## RESULTS

### Urp1 and Urp2 peptides are dispensable for embryonic axial straightening

To determine whether Urp1 and Urp2 are required for spine morphology, we used CRISPR/Cas9 to generate zebrafish mutant lines. We used pairs of guide RNAs to induce deletions across the genetic region coding for the 10-amino acid secreted peptides (Fig. 1A-B and S1A-D). This resulted in mutant lines that we refer to as *urp1*^*ΔP*^ and *urp2*^*ΔP*^ because they lack the peptide coding sequence. In addition, mRNA quantitation revealed downregulation of *urp1* and *urp2* in their respective mutant backgrounds, indicating transcript decay (Fig. S1E).

**Figure 1.**
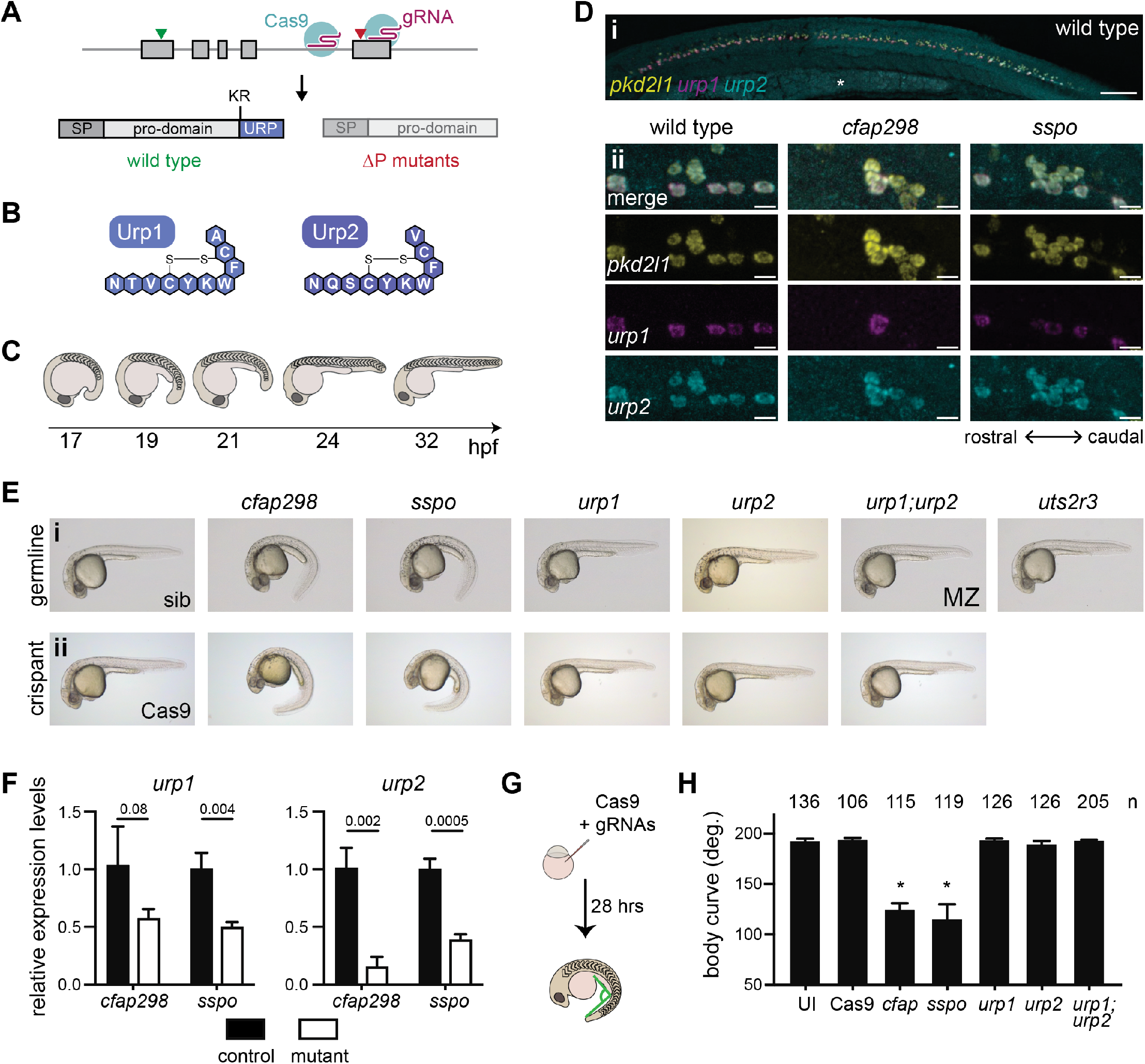
Urp1 and Urp2 are dispensable for axial straightening. **A)** *urp1* and *urp2* are 5-exon genes (gray boxes). The final exon codes for the 10-amino acid peptides, produced after cleavage from the prodomain at a dibasic site (KR). Pairs of gRNAs were used to induce deletions of Urp1 and Urp2 peptide coding sequences in *urp1*^*ΔP*^ and *urp2*^*ΔP*^ mutants, respectively. SP – signal peptide. **B)** Urp1 and Urp2 peptide sequences with identical hexacyclic regions. **C)** Zebrafish posterior axial straightening, the morphogenetic process which straightens the embryonic body. **D)** Fluorescence *in situ* hybridization based on hybridization chain reaction (*in situ* HCR) analysis of *pkd2l1, urp1* and *urp2* expression in the central canal at 28 hpf. *pkd2l1* expression marks CSF-cNs. *urp1* expression is restricted to dorsal CSF-cNs while *urp2* is expression in all CSF-cNs. Both *urp1* and *urp2* are expressed in *cfap298*^*tm304*^ and *sspo*^*b1446*^ mutants, though comparison of expression between samples was non-quantitative. (i) Shows the zebrafish trunk with the yolk stalk labelled (*). (ii) Shows zoomed regions taken at the rostro-caudal level at the end of the yolk stalk. Scale bars: 150 µm (i), 10 µm (ii). **E)** Lateral views of 28-30 hpf germline mutants (i) and crispants (ii). The *urp1*^*ΔP*^*;urp2*^*ΔP*^ double mutants are maternal zygotic (MZ) mutants. Sibling (sib) and Cas9-only injected embryos served as controls. **F)** Quantitative reverse transcriptase PCR (qRT-PCR) analysis of *urp1* and *urp2* mRNA expression levels in *cfap298*^*tm304*^ and *sspo*^*b1446*^ mutants at 28 hpf. *n* > 3 biologically independent samples. Bars represent mean ± s.e.m. Two-tailed student’s *t* test used to calculate *P* values. **G)** Schematic of crispant generation and body curve analysis. **H)** Quantitation of crispant body curves where bars represent mean ± s.d. for at least three independent clutches and injection mixes. The total number of embryos analyzed is given. \**P* < 0.0001, student’s *t* test applied. UI – uninjected.

We first assessed *urp1*^*ΔP*^ and *urp2*^*ΔP*^ mutants for embryonic phenotypes. A previous morpholino-based knockdown study concluded that Urp1 and Urp2 are required for axial straightening, the process by which the ventrally curved zebrafish embryo straightens as the trunk elongates and detaches from the yolk (Fig. 1C). Urp1/Urp2 morphants failed to undergo straightening and therefore displayed CTD (Zhang et al., 2018). Surprisingly, both *urp1*^*ΔP*^ and *urp2*^*ΔP*^ mutants underwent normal axial straightening and did not exhibit CTD (Fig. 1Ei). By contrast, we observed CTD in both *cfap298*^*tm304*^ mutants that lack cilia motility in the central canal (Bearce et al., 2022) and *sspo*^*b1446*^ mutants in which the RF constituent SCOspondin is mutated, as expected (Fig. 1Ei and S2A-B). Notably, *cfap298*^*tm304*^ and *sspo*^*b1446*^ mutants maintained *urp1* and *urp2* expression in CSF-cNs, central canal neurons marked by *pkd2l1* expression (Fig. 1D), though *urp1* and *urp2* transcripts were quantitatively reduced (Fig. 1F). We reasoned that the absence of CTD in *urp1*^*ΔP*^ and *urp2*^*ΔP*^ mutants might reflect redundancy, since Urp1 and Urp2 peptides are highly similar, with identical hexacyclic regions (Fig. 1B and S1A-B). Alternatively, maternally-derived *urp1* and/ or *urp2* transcripts may function to prevent phenotypes from manifesting. However, maternal zygotic (MZ) *urp1*^*ΔP*^;*urp2*^*ΔP*^ double mutants also exhibited linear body axes (Fig. 1Ei), ruling out redundant or maternal gene product function. This demonstrates that Urp1 and Urp2 peptide-null mutants undergo axial straightening.

To confirm this finding, we performed additional Urp1 and Urp2 loss of function experiments. By injecting multiple guide RNAs along with Cas9 into wild-type embryos at the one-cell stage, we generated mosaic mutants, called crispants, that were then screened for body shape phenotypes (Fig. 1G). In positive control experiments, *cfap298* and *sspo* crispants exhibited robust CTD, phenocopying germline *cfap298*^*tm304*^ and *sspo*^*b1446*^ mutants (Fig. 1Eii). Quantitation of body curvature revealed that crispant generation was highly efficient, with CTD penetrance being close to 100% (Fig. 1H). By contrast, *urp1* and *urp2* single and double crispants exhibited straight body axes that were not different to uninjected embryos or embryos injected with Cas9 only (Fig. 1Eii and H). We used the AB genetic background for the majority of our work but we also generated and phenotyped *urp1;urp2* double crispants on WIK and TU backgrounds to test for potential background effects. Normal axial straightening upon mutation of *urp1* and *urp2* was also observed on these backgrounds (Fig. S1F). Overall, crispant results confirmed germline mutant findings. We conclude that Urp1 and Urp2 peptides are dispensable for axial straightening in embryonic zebrafish.

### Urp1 and Urp2 function redundantly to maintain spine morphology

Next, we determined the impact of Urp1 and Urp2 loss on adult spine morphology. Outwardly, *urp1*^*ΔP*^ mutant adults appeared normal whereas *urp2*^*ΔP*^ mutants exhibited minor body dysmorphologies and kinked tails (n=72 for *urp1*^*ΔP*^ mutants and n=92 for *urp1*^*ΔP*^ mutants, Fig. S3A). To assess spine morphology directly, we imaged bone by X-ray microcomputed tomography (µCT) scanning. Three-dimensional reconstitutions of µCT data from 3-month-old fish showed that *urp1*^*ΔP*^ mutants indeed exhibited overtly normal skeletal morphology (n=7) while *urp2*^*ΔP*^ mutants showed slight sagittal curves (n=4; Fig. 2A-C, S3A and Movies S1-3). By contrast to these absent or mild deformities in single mutants, *urp1*^*ΔP*^*;urp2*^*ΔP*^ double mutants exhibited prominent curves, with significant dorsal-ventral Cobb angles, a measure of deviation from straightness, especially in the caudal region of the spine (Fig. 2D, 2F-G, 2H, S3A-B and Movie S4). These data indicate that Urp1 and Urp2 are essential for adult spine morphology and that they function in a mostly redundant fashion in this context.

**Figure 2.**
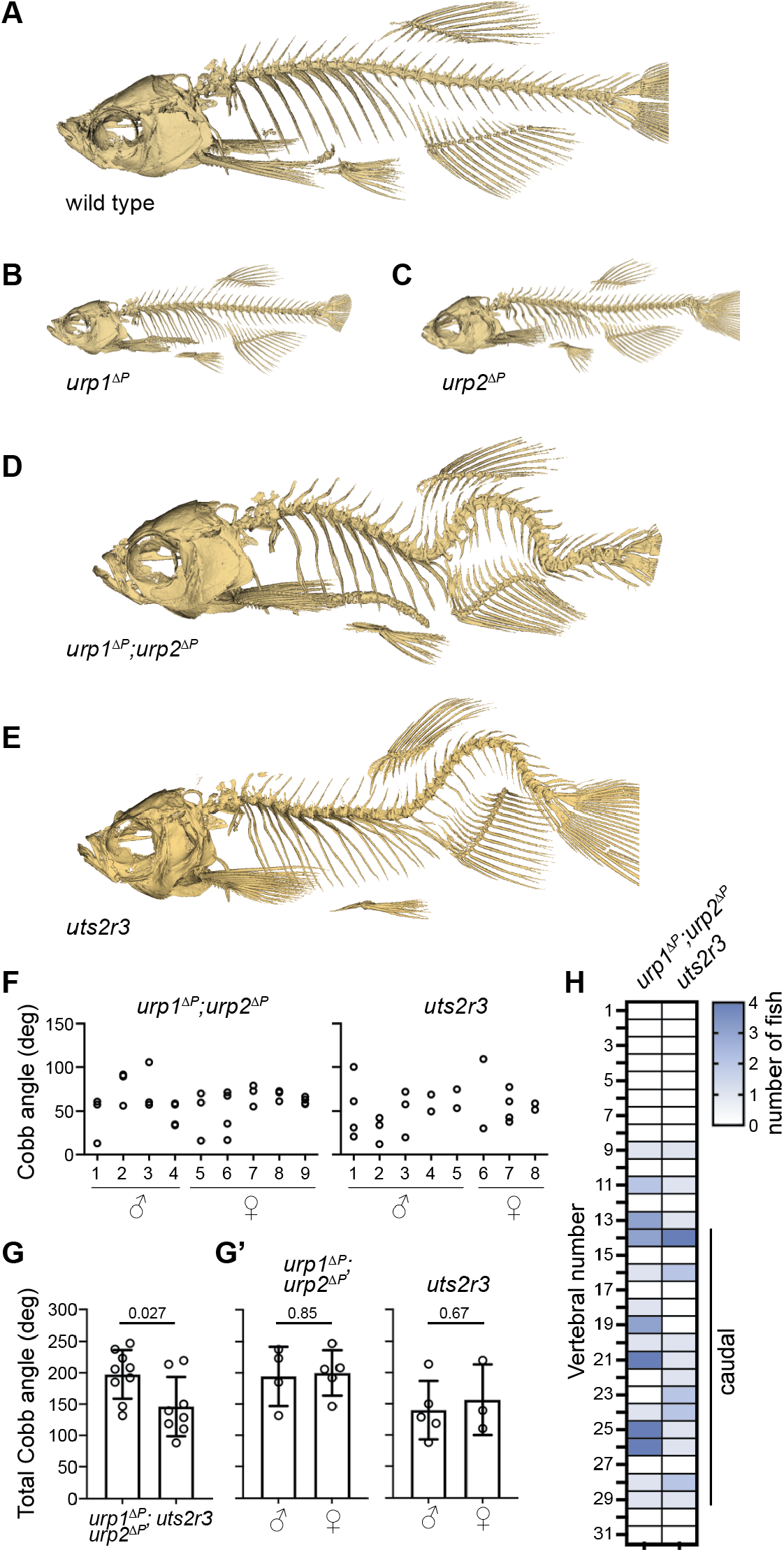
Urp1 and Urp2 are required for proper adult spine morphology. **A-E)** Lateral views of µCT reconstitutions of wild-type (A), and *urp1*^*ΔP*^ (B), *urp2*^*ΔP*^ (C), *urp1*^*ΔP*^*;urp2*^*ΔP*^ (D) and *uts2r3*^*b1436*^ (E) mutants at 3-months of age. **F)** Cobb angle measurements for individual fish in the sagittal plane for *urp1*^*ΔP*^*;urp2*^*ΔP*^ and *uts2r3*^*b1436*^ mutants. Circles represent angles for individual curves. **G-G’)** Total Cobb angles with each circle representing an individual fish. The mean ± s.d. is shown. G’ is the data from G parsed for sex. *P* values are given from two-tailed unpaired Student’s *t*-tests. **H)** The position of curve apex is plotted and shows that most curves are in caudal vertebrae.

### Urp1 and Urp2 signal through the Uts2r3 receptor to control spine morphology

Urp1 and Urp2 peptides engage G-protein coupled receptors (Ames et al., 1999; Chatenet et al., 2004; Elshourbagy et al., 2002; Labarrere et al., 2003; Liu et al., 1999; Nothacker et al., 1999). While a single Urotensin II receptor (UT) gene is found in humans, and has recently been linked to abnormal spinal curvature (Dai et al., 2021), the zebrafish genome encodes five such receptors. One of those, Uts2r3, was previously implicated in spine morphology (Zhang et al., 2018). To systematically compare the effects of Uts2r3 receptor mutation with loss of Urp1 and Urp2, we generated a *uts2r3* mutant line harboring a 178-amino acid deletion after the third amino acid, significantly disrupting the protein (Fig. S2C). Like *urp1*^*ΔP*^*;urp2*^*ΔP*^ double mutants, these *uts2r3*^*b1436*^ mutants underwent normal axial straightening as embryos (Fig. 1Ei) and went on to exhibit spinal curves as adults (Fig. 2E, S3A and Movie S5). Cobb angle measurements showed that *uts2r3*^*b1436*^ mutants and *urp1*^*ΔP*^*;urp2*^*ΔP*^ mutants were similar, though curves in *urp1*^*ΔP*^*;urp2*^*ΔP*^ mutants were slightly more severe (Fig. 2F and G). As for *urp1*^*ΔP*^*;urp2*^*ΔP*^ mutants, *uts2r3*^*b1436*^ mutants showed mostly caudally-located curves, especially in the most rostral of the caudal vertebrae (Fig. 2H). Thus, although we cannot rule out minor roles for other UT receptors, these data suggest that Urp1 and Urp2 control spine morphology largely by signaling through Uts2r3.

### Urotensin pathway mutants display adolescent-onset spinal curves in the absence of structural vertebral defects

Next, we determined whether Urotensin pathway mutants recapitulated any signs of disease present in patients. Several types of human spinal curves onset during adolescent growth (Cheng et al., 2015). To discern the stage of onset of curves in Urotensin pathway mutants, we monitored *urp1*^*ΔP*^*;urp2*^*ΔP*^ double mutant cohorts as they grew. Subtle curves first became apparent between 9-11 dpf, corresponding to a standard length between 3.9 ± 0.7 mm and 5.9 ± 0.4 mm (mean ± s.d.; Fig. 3A-B and Movie S6), a stage when adolescents were rapidly growing (Fig. 3C). By 13 dpf (standard length 6.2 ± 0.3 mm), curves were evident in all *urp1*^*ΔP*^*;urp2*^*ΔP*^ mutants and progressively worsened up to 17 dpf (8.3 ± 0.4 mm) when we ended this analysis (Fig. 3A-B). At 30 dpf, we assessed *urp1*^*ΔP*^*;urp2*^*ΔP*^ mutants by µCT and found variability in curve position and amplitude (Fig. 3D and S3C). Notably, at this stage, several mutants exhibited a significant curve in the pre-caudal vertebrae, in addition to a caudal curve (Fig. 3D and S3C). Since pre-caudal curves were rare in mutants at 3-months, this suggested that curve location is dynamic and that pre-caudal curves form then resolve in some mutants as they grow to adulthood.

**Figure 3.**
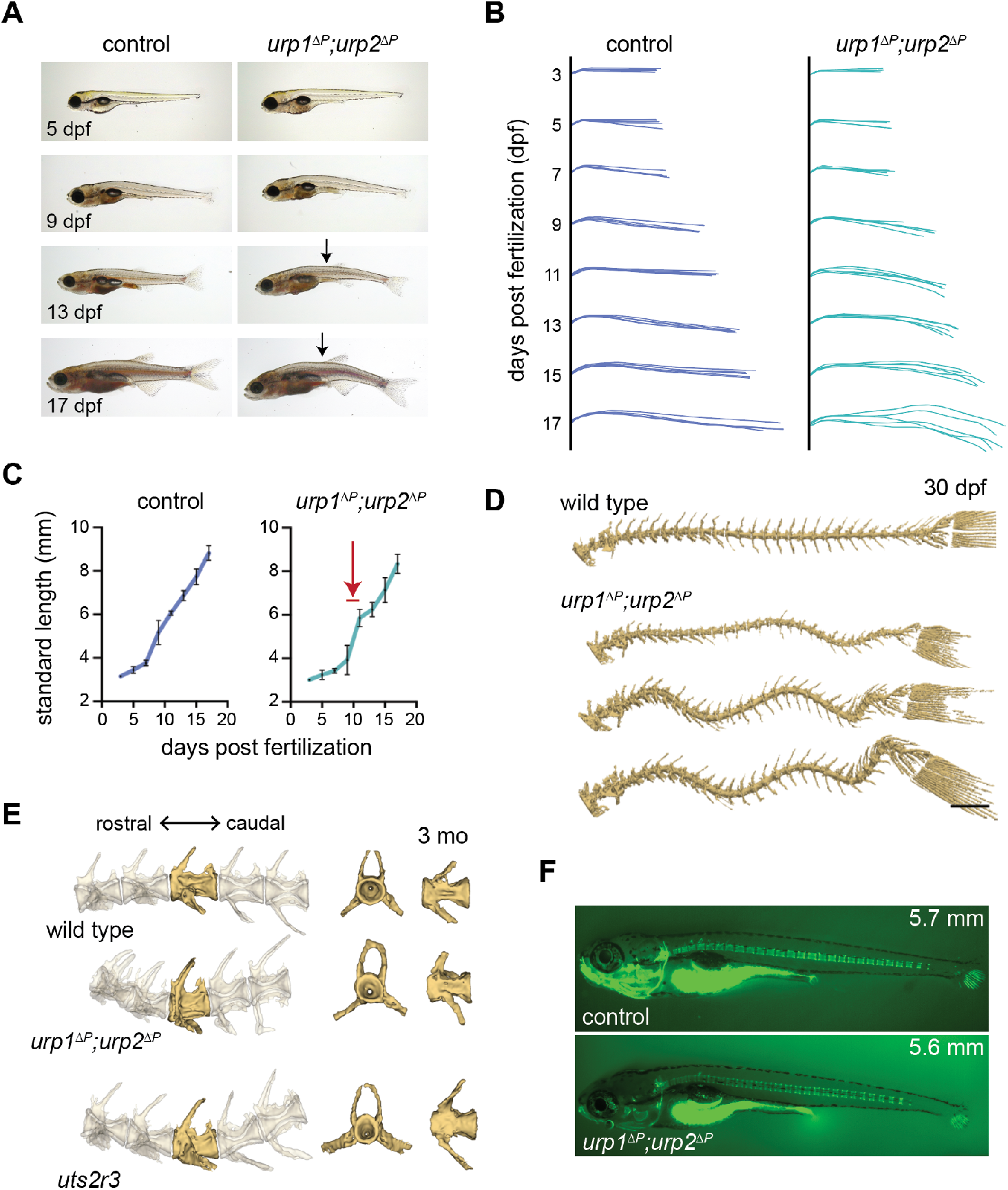
*urp1*^*ΔP*^*;urp2*^*ΔP*^ mutants exhibit adolescent-onset spinal curves without structural vertebral defects. **A)** Lateral views of control fish and age-matched *urp1*^*ΔP*^*;urp2*^*ΔP*^ mutants. Arrows point to forming body curves. **B)** Traces of body shape every 2 days for 5 fish per time point from 3-17 dpf. **C)** Growth curves for control and *urp1*^*ΔP*^*;urp2*^*ΔP*^ mutants were indistinguishable. Arrow shows time of curve onset. **D)** µCT reconstitutions of spines at 30 dpf with heads, fins and ribs removed. Scale bar: 1 mm **E)** µCT reconstitutions of three pre-caudal and two caudal vertebrae including frontal and lateral views of the highlighted vertebra. No structural defects were observed in *urp1*^*ΔP*^*;urp2*^*ΔP*^ or *urs2r3*^*b1436*^ mutants. **F)** Calcein staining revealed well-structured vertebrae forming in controls and *urp1*^*ΔP*^*;urp2*^*ΔP*^ mutants, with fish standard length given in mm.

Next, we assessed whether spinal curves in Urotensin pathway mutants were caused by congenital-like defects in vertebral patterning or structure. Careful analysis of µCT datasets as well as staining of juveniles with the vital dye calcein, revealed no defects in vertebral patterning or overt structure in *urp1*^*ΔP*^*;urp2*^*ΔP*^ and *uts2r3*^*b1436*^ mutants (Figs. 3E-F and Movie S7). This demonstrates that adolescent-onset curves in Urotensin-deficient mutants occur in the absence of vertebral patterning or structural defects. Additionally, we parsed our phenotypic data for sex since spinal curves often show sex bias in severity in humans (Cheng et al., 2015), something which has been recapitulated in some zebrafish spinal curve models (Marie-Hardy et al., 2021). However, in both *urp1*^*ΔP*^*;urp2*^*ΔP*^ and *uts2r3*^*b1436*^ mutants, we found no significant differences in curve penetrance or severity between males and females (Figs. 2F and 2G’).

### Urotensin-deficient mutants are phenotypically distinct from *cfap298*^*tm304*^

We next compared the phenotypes of Urotensin pathway mutants to other mutant lines that exhibit spinal curves. The *cfap298*^*tm304*^ line harbors a temperature-sensitive mutation in *cfap298*, a gene required for cilia motility in several organisms including humans (Austin-Tse et al., 2013; Bearce et al., 2022; Jaffe et al., 2016). *cfap298*^*tm304*^ mutants exhibit reduced cilia motility in the central canal and, if the resulting CTD is embryonically rescued by temperature shifts, develop adolescent-onset spinal curves (Grimes et al., 2016).

These curves were argued to model an adolescent idiopathic scoliosis (AIS)-like condition (Fig. 4A-B and Movie S8; Grimes et al., 2016; Marie-Hardy et al., 2021). Importantly, both *urp1* and *urp2* transcripts were significantly downregulated in *cfap298*^*tm304*^ mutants (Fig. 1F) suggesting that spinal curves in *cfap298*^*tm304*^ might be the result of reduced Urp1/Urp2 expression. To systematically compare *cfap298*^*tm304*^ mutants with Urotensin-deficient mutants, we raised cohorts of *urp1*^*ΔP*^*;urp2*^*ΔP*^ mutants, *uts2r3*^*b1436*^ mutants and temperature-shift-rescued *cfap298*^*tm304*^ mutants alongside one another in the same aquatics facility after backcrossing all lines to the AB strain for multiple generations. At 3-months, we performed µCT scanning and three-dimensional reconstitutions. First, we calculated dorsal-ventral Cobb angles, which revealed that *cfap298*^*tm304*^ mutants were more severely curved (average total Cobb angle: 247.4 ± 23.7 degrees) than either *urp1*^*ΔP*^*;urp2*^*ΔP*^ mutants (197.5 ± 38.9 degrees) or *uts2r3*^*b1436*^ mutants (146.1 ± 47.4 degrees) (compare Figs. 4C-D with 2F-G’). Second, *cfap298*^*tm304*^ mutants showed prominent dorsal-ventral curves in the precaudal as well as caudal vertebrae, a distinct pattern compared with the predominantly caudal curves in *urp1*^*ΔP*^*;urp2*^*ΔP*^ and *uts2r3*^*b1436*^ mutants (compare Figs. 4A-B with 2D-E and Fig. S3C). Third, *cfap298*^*tm304*^ mutants exhibited significant lateral curvature of the spine, often with spinal twisting, a hallmark of AIS-like curves (Fig. 4E). By contrast, *urp1*^*ΔP*^*;urp2*^*ΔP*^ (Fig. 4E) and *uts2r3*^*b1436*^ mutants (not shown) showed planar curves, with very minor or no lateral deviations (Fig. 4E). These results demonstrated that cilia motility mutants and Urotensin-deficient mutants exhibit distinct spinal curve phenotypes. As such, the causes of spinal curves in *cfap298*^*tm304*^ mutants can only be partially explained by reduced Urp1/Urp2 expression.

**Figure 4.**
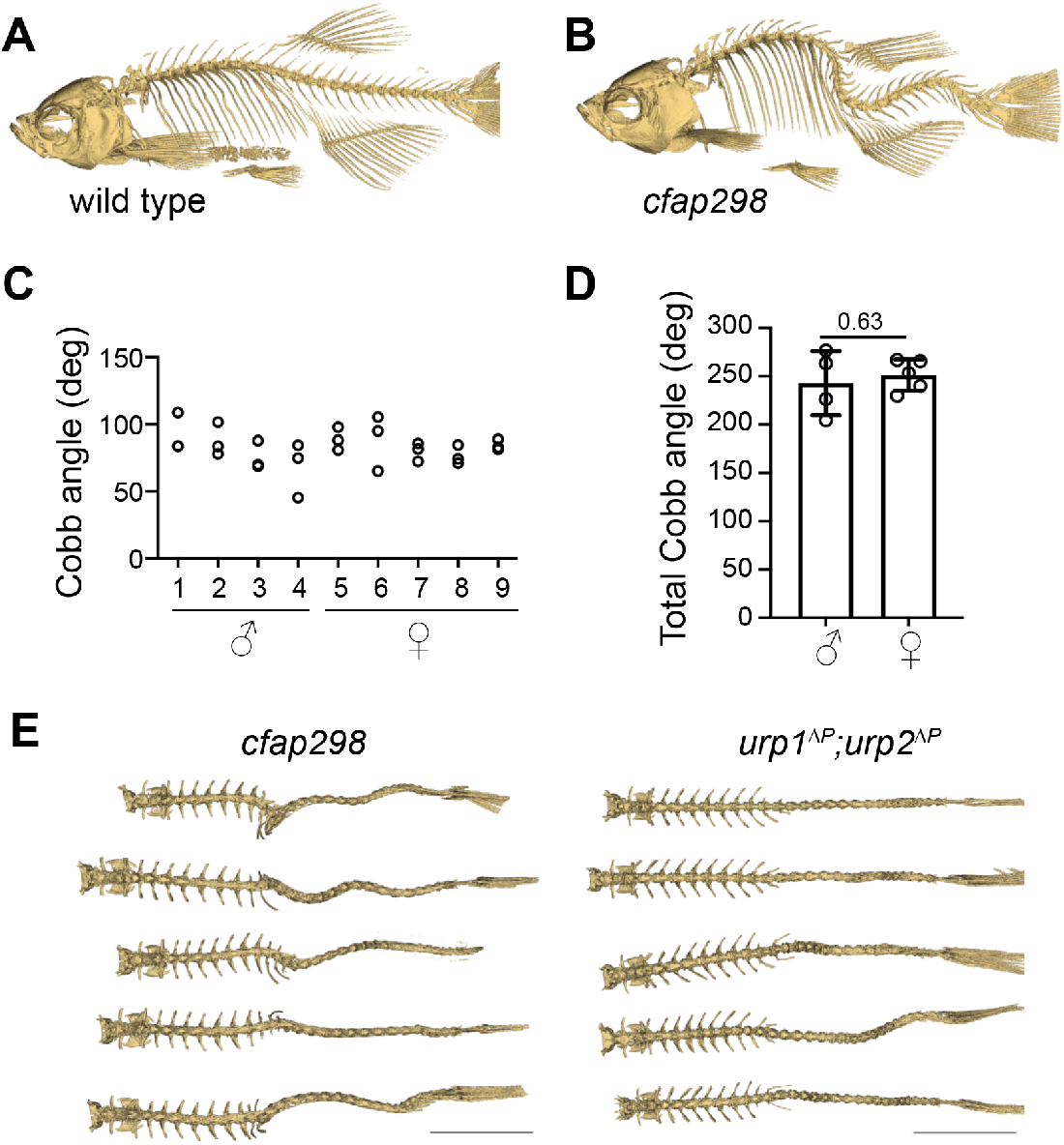
*urp1*^*ΔP*^*;urp2*^*ΔP*^ mutants and *cfap298*^*tm304*^ mutants are phenotypically distinct. **A-B)** Lateral views of µCT reconstitutions of wild-type (A), and *cfap298*^*tm304*^ mutants (B). **C)** Cobb angle measurements for individual fish in the sagittal plane for *cfap298*^*tm304*^ mutants. Circles represent angles for individual curves. **D)** Total Cobb angles with each circle representing an individual fish. The mean ± s.d. is shown. *P* values are given from two-tailed unpaired Student’s *t*-tests. **E)** Dorsal views of µCT reconstitutions with ribs and fins removed. Scale bars: 5 mm.

Pkd2l1 is a Polycystin family ion channel expressed in CSF-cNs (Sternberg et al., 2018), the same cell type which expresses Urp1 and Urp2 (Fig. 1D; Quan et al., 2015). Pkd2l1 is responsible for flow-induced Ca^2+^ signaling in CSF-cNs (Böhm et al., 2016; Sternberg et al., 2018). While *pkd2l1*^*icm02*^ mutants exhibited normal early axis development they go on to develop mild kyphosis-like curves upon aging (Sternberg et al., 2018), a result we recapitulated after raising *pkd2l1*^*icm02*^ mutants on the same genetic background and under the same conditions as Urotensin-deficient mutants for direct comparison (data not shown). This kyphosis-like phenotype was again highly distinct from *urp1*^*ΔP*^*;urp2*^*ΔP*^ and *uts2r3*^*b1436*^ mutants.

Overall, our phenotypic data show that Urotensin pathway mutants develop adolescent-onset curves without vertebral structural defects or sex bias. Coupled to the consistently caudal location of curves in adults as well as the lack of lateral deviation, we suggest that Urotensin pathway mutants most closely reflect a lordosis-like condition. In agreement, Urotensin mutants were phenotypically distinct from established AIS (*cfap298*^*tm304*^) and kyphosis (*pkd2l1*^*icm02*^) models.

### RF breakdown precedes AIS-like curves in cfap298^tm304^ mutants

Given the links between motile cilia, the RF and *urp1* and *urp2* expression (Fig. 1D and 1F; Cantaut-Belarif et al., 2020; Liu et al., 2020; Zhang et al., 2018) as well as the requirement for proper RF assembly to prevent spinal curves (Rose et al., 2020; Troutwine et al., 2020), we assessed Sspo, the major component of the RF, in spinal curve mutants. To visualize Sspo localization, we used the *sspo-GFP*^*ut24*^ line in which GFP is fused to the endogenous *sspo* locus (Troutwine et al., 2020). First, we assessed Sspo localization in the central canal of *cfap298*^*tm304*^ mutants at 28 hpf. In sibling controls, Sspo localized into an RF throughout the central canal (Fig. 5A). By contrast, *cfap298*^*tm304*^ mutants raised at restrictive temperatures, which exhibited reduced central canal cilia motility and CTD (Fig. 1E; Bearce et al., 2022), lacked RF. Instead, Sspo was diffusely localized in the central canal in these mutants (Fig. 5B), in agreement with previous work showing that cilia motility is required for RF assembly in embryos (Cantaut-Belarif et al., 2018).

**Figure 5.**
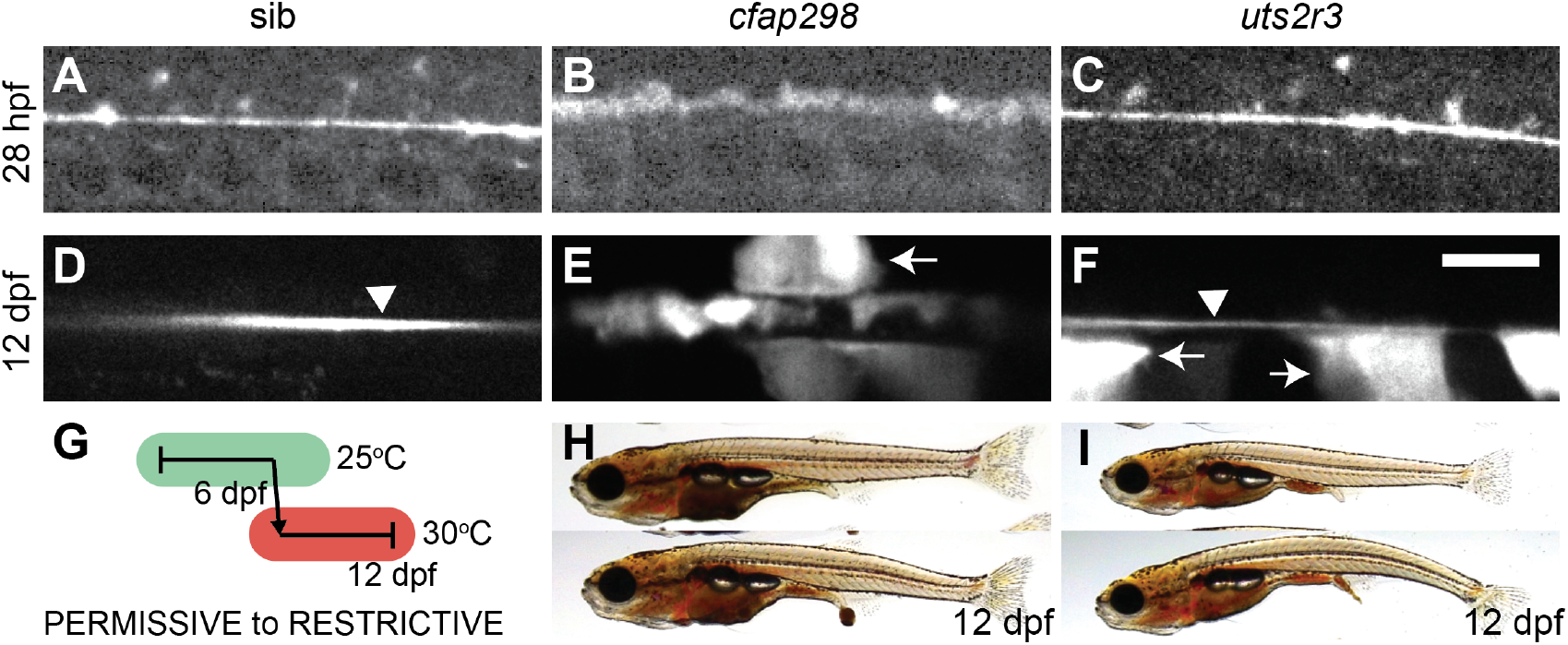
RF breakdown in *cfap298*^*tm304*^ mutants but not Urotensin-deficient mutants. **A-F)** Grayscale maximal intensity projection of Sspo-GFP localization in the central canal in 28 hpf embryos (A-C) and 12 dpf adolescents (D-F). RF is denoted by arrow heads in D and F. Arrows point to structures along the central canal that become GFP-positive in *cfap298*^*tm304*^ and *uts2r3*^*b1436*^ mutants. Scale bar: 10 µm. **G)** Schematic of temperature shift experiment in which *cfap298*^*tm304*^ mutants are initially raised at permissive temperatures before being shifted to restrictive temperatures at 6 dpf, then imaged at 12 dpf. **H-I)** Lateral views of *cfap298*^*tm304*^ (H) and *uts2r3*^*b1436*^ (I) mutants at 12 dpf when Sspo-GFP imaging took place.

Next, we took advantage of the temperature-sensitive nature of the *cfap298*^*tm304*^ mutation to determine whether RF is also disrupted at later timepoints, during adolescent stages when AIS-like curves begin to develop. To do so, we initially raised *cfap298*^*tm304*^ mutants at permissive temperatures, allowing RF to correctly form and embryos to fully straighten. At 6 dpf, we transitioned larvae to restrictive temperatures (Fig. 5G), something which led to the appearance of curves by 11-14 dpf (standard length 6.5 ± 0.2 mm), then assessed Sspo localization at 12 dpf. As in embryos, Sspo localized into a defined RF in the central canal in sibling controls (arrow head in Fig. 5D) but was diffuse in temperature upshifted *cfap298*^*tm304*^ mutants, both in mutants that had yet to develop curves and those exhibiting subtle curves (Fig. 5E and 5H). This demonstrates 1) that cilia motility is not only required for the initial formation of the RF but also for its maintenance; and 2) breakdown of the RF precedes curve onset in cilia motility-deficient mutants. This supports a model in which continued cilia motility maintains the RF structure and loss of the RF causes AIS-like curves to develop in cilia motility mutants.

### The RF remains intact in Urotensin-deficient mutants both before and after curve formation

Next, we imaged Sspo-GFP localization in the central canal of *uts2r3*^*b1436*^ mutants to determine whether RF breakdown could be causative in Urotensin-associated spinal curves. In agreement with that lack of axial straightening phenotype in *uts2r3*^*b1436*^ mutant embryos (Fig. 1Ei), we found that Sspo localized normally into the RF in the central canal at 28 hpf in *uts2r3*^*b1436*^ mutants (Fig. 5C). At 12 dpf, as spinal curves were beginning to manifest (Fig. 5I), Sspo still formed an intact RF in *uts2r3*^*b1436*^ mutants (arrow head in Fig. 5F). The fact that RF is present in *uts2r3*^*b1436*^ mutants as curves form suggests that Urotensin-associated curves are not caused by defective RF formation. This result coheres with the distinct spinal phenotypes exhibited by *cfap298*^*tm304*^ mutants and *uts2r3*^*b1436*^ mutants.

While imaging Sspo-GFP at 12 dpf, we noted that in both *cfap298*^*tm304*^ mutants and *uts2r3*^*b1436*^ mutants, large central canal cells with the appearance of CSF-cNs became GFP-positive (arrows in Figs. 5E-F), something we rarely observed in control fish (Fig. 5D). Indeed, the GFP-positive central canal cells in *uts2r3*^*b1436*^ give the effect of making the RF appear comparatively smaller/dimmer (compare RF in Fig. 5F and 5D). We suggest that CSF-cNs may endocytose Sspo-GFP monomers in *cfap298*^*tm304*^ mutants where the RF has broken down and in *uts2r3*^*b1436*^ mutants where RF is likely to be making increasing numbers of contacts with CSF-cNs (Orts-Del’Immagine et al., 2020) owing to the onset of spinal curves.

## DISCUSSION

Urotensin II (UII) is a cyclic peptide that was first identified from the teleost urophysis (Pearson et al., 1980) and subsequently found to exist in amphibians (Conlon et al., 1992) and mammals (Coulouarn et al., 1998). A highly similar peptide, called Urotensin II-related peptide (URP) was then isolated from the brains of rodents (Sugo et al., 2003). UII and URP both signal via the Urotensin receptor (UT), a G protein-coupled receptor (Ames et al., 1999; Liu et al., 1999; Mori et al., 1999; Nothacker et al., 1999). UII’s, URPs and UTs have been linked to cardiovascular function and inflammation but their roles in the development of morphology are little understood.

In this study, we discovered a role for two of the URPs, Urp1 and Urp2, in zebrafish spine morphology. To do so, we generated *urp1*^*ΔP*^ and *urp2*^*ΔP*^ mutants that lacked the genetic region coding for the Urp1 and Urp2 peptides, respectively, then phenotyped skeletal morphology by µCT. This revealed that Urp1 and Urp2 function redundantly to control spine morphology, with double mutants, as well as Uts2r3 receptor mutants, developing spinal curves in the caudal region during adolescent growth. The lack of vertebral defects, the location and direction of the curves coupled with phenotypic differences compared with mutants that model AIS and kyphosis, suggested that Urotensin-deficient mutants model a lordosis-like condition.

The RF, a long proteinaceous thread-like structure which sits in the central canal, has been implicated in controlling body axis and spine morphology (Cantaut-Belarif et al., 2018; Lu et al., 2020; Rose et al., 2020; Troutwine et al., 2020). We find that RF breaks down prior to curve formation in the cilia motility *cfap298*^*tm304*^ mutant. Given other studies linking presence of the RF with a linear body axis, this strongly suggests that RF breakdown causes spinal curves in *cfap298*^*tm304*^ mutants. By contrast, Urotensin deficient mutants exhibited an intact RF, demonstrating that curves are not formed by RF breakdown in Urotensin mutants and nor do the presence of curves significantly disrupt RF structure. This coheres with a model in which Urotensin signals act downstream of RF function in controlling spine morphology. Similarly, *urp1* and *urp2* expression is known to be controlled by RF function during embryonic phases (Fig. 1E; Cantaut-Belarif et al., 2020; Lu et al., 2020; Rose et al., 2020; Zhang et al., 2018)

Intriguingly, motile cilia mutants and RF mutants exhibit three-dimensional spinal curves, with dorsoventral and medio-lateral curvature (Grimes et al., 2016; Lu et al., 2020; Rose et al., 2020; Troutwine et al., 2020). By contrast, we find that Urotensin-deficient mutants exhibit largely planar curves, only in the dorso-ventral direction. Urp1 and Urp2 are expressed in CSF-cNs (Fig. 1D; Quan et al., 2015), a cell type which consists of both dorsal and ventral subpopulations. While Urp1 and Urp2 are co-expressed in ventral CSF-cNs, only Urp2 is expressed in dorsal CSF-cNs (Fig. 1D; Quan et al., 2015). It is therefore tempting to speculate that dorso-ventral spine shape is mediated specifically by ventral CSF-cNs which express higher amounts of Urp1/Urp2 peptides, resulting in dorso-ventral curves in Urotensin-deficient mutants. This also suggests, as above, that while Urp1/ Urp2 expression is in part controlled by upstream cilia motility and RF function, decreased Urotensin signaling cannot account fully for the spinal curve phenotypes that occur upon loss of cilia motility or the RF. Indeed, the RF is required for both dorsal *and* ventral CSF-cN function (Orts-Del’Immagine et al., 2020), which may explain why dorso-ventral and medio-lateral curves occur when RF is disrupted either by cilia motility mutations or mutations to SCOspondin. This scenario is further complicated by the finding that increased *urp1* and *urp2* expression occurs upon mutation of *rpgrip1l*, a gene encoding a component of the ciliary transition zone (Vesque et al., 2019 — preprint). As such, various cilia-dependent signals likely control the precise levels of Urp1/Urp2 peptides, and controlling those levels during important growth stages appears critical for maintaining the shape of the spine.

A surprising finding from our work was that Urp1 and Urp2 peptides are genetically dispensable for embryonic axial straightening. This interpretation is challenged by morpholino knockdown of Urp1/Urp2, which does result in failure of axial straightening in some individuals, resulting in a curly tail down (CTD) phenotype (Zhang et al., 2018). One possibility is that the CTD of morphants results from morpholino off-target effects. However, this seems unlikely for three reasons: 1) adding exogenous Urp1/Urp2 peptides to the central canal can rescue the CTD phenotype of cilia motility- and RF-deficient mutants (Lu et al., 2020; Zhang et al., 2018), suggesting the involvement of Urp1/Urp2 in axial straightening, at least in gain-of-function experiments; 2) since motile cilia and the RF are required for both axial straightening during embryogenesis and for the maintenance of spine morphology during adolescence, it seems parsimonious that Urp1/Urp2 peptides would also function across these two life stages; and 3) *urp1* and *urp2* transcript levels are reduced in motile cilia and RF mutants (Fig. 1F; Cantaut-Belarif et al., 2020; Lu et al., 2020; Zhang et al., 2018), suggestive of a link between Urp1 and Urp2 upregulation and axial straightening.

Nevertheless, the lack of CTD in our mutants, in which the Urp1 and Urp2 peptide coding sequences were entirely removed, is clear: single, double and maternal-zygotic *urp1*^*ΔP*^ and *urp2*^*ΔP*^ mutants all underwent normal axial straightening. This strongly argues that Urp1 and Urp2 are dispensable for straightening. Since the deletions were induced towards the end of the protein, it does leave open the possibility that the pro-domain sequences are required for straightening. However, this seems unlikely for three reasons: 1) *urp1*^*ΔP*^ and *urp2*^*ΔP*^ mutants showed *urp1* and *urp2* transcript downregulation, respectively, in addition to deletion of the peptide coding regions; 2) *urp1* and *urp2* single and double crispants, in which gRNAs targeted several regions of the gene, also showed normal straightening; and 3) exogenous addition of Urp1 and Urp2 peptides, without pro-domains, rescued CTD of a cilia motility and RF mutant (Lu et al., 2020; Zhang et al., 2018), suggesting that it is the peptide itself and not some other region which is functional.

One possibility is that genetic compensation explains the mutant/morphant phenotypic differences (Rossi et al., 2015; Sztal and Stainier, 2020). In this putative scenario, a feedback response in the cell buffers otherwise harmful mutations, preventing their effects from manifesting phenotypically. A recently discovered compensation mechanism is transcriptional adaptation in which mutant mRNA is decayed and decay products are, via a sequence-dependent mechanism, recruited to genes with similar sequences where they promote transcriptional upregulation (El-Brolosy et al., 2019; Ma et al., 2019). The upregulation of adapting genes masks phenotypes in mutants but not morphants. We did find slight upregulation of *urp2* in *urp1*^*ΔP*^ mutants (Fig. S1E), which may indicate transcriptional adaptation, although this alone cannot explain the lack of CTD phenotypes in *urp1*^*ΔP*^ mutants because *urp1*^*ΔP*^*;urp2*^*ΔP*^ double mutants also lack CTD. It will be informative in the future to determine whether other Urotensin II-encoding peptides (Fig. S1A-B) are able to compensate, during embryonic phases, for loss of *urp1* and *urp2* or if other factors explain the mutant/ morphant discrepancies.

While we were in the final stages of preparing the current manuscript, a study was released which made several complementary findings, also concluding that Urp1 and Urp2 function redundantly to maintain spine shape (Gaillard et al., 2022 — preprint). Importantly, and in agreement with our work, Gaillard and colleagues observed no embryonic axial defects upon genetic loss of Urp1 and Urp2. Moreover, they found that other Urotensin II-encoding genes were not upregulated in *urp1* and *urp2* mutants, arguing against phenotypic masking by genetic compensation. Our findings and those of Gaillard and colleagues together strongly argue that Urp1 and Urp2 are not essential for axial straightening during embryogenesis, but are instead required for the maintenance of the body axis during growth. As such, other mechanisms, currently unknown, likely operate downstream of cilia motility and RF function to mediate embryonic axial straightening.

Future efforts will be required to discern which tissues respond to Urp1/Urp2 signals during the control of spine shape. During embryonic stages, Uts2r3 is expressed in dorsal muscle, but it remains unclear how Urp1/Urp2 peptides released by CSF-cNs could signal to effect muscle during adolescent stages. Indeed, inhibition of acetylcholine receptors to prevent neuromuscular signaling does not prevent axial straightening. Moreover, based on single cell RNA-sequencing gene expression atlases (Farnsworth et al., 2019), *uts2r3* is expressed in several other cell types in addition to muscle. Tissue-specific ablations and rescue experiments should be used to untangle precisely where Uts2r3-dependent Urp1/Urp2 signaling occurs to control spine morphology. It will also be critical to determine the timing of action of Urotensin signaling. While we observe phenotypes first appearing between 9-11 dpf, it is possible that the underlying defect in mutants is caused by an earlier event which only phenotypically manifests later. One candidate is subtle disruptions to the notochord, which may later result in spinal curves.

A major question is whether the function of Urotensin signaling in spine morphology is conserved in other species and whether our findings are directly relevant to humans. These questions will require future work to answer but two recent studies shed some light on these matters. First, in the frog *Xenopus laevis*, disruption of Utr4, a counterpart of Uts2r3, causes abnormal curvature of the body axis (Alejevski et al., 2021). Second, a human genetics study reported that rare mutations in UTS2R are significantly associated with spinal curvature, being discovered within AIS patient cohorts (Dai et al., 2021). Thus, a deeper understanding of the role of Urotensin signaling in maintaining spinal shape will not only provide insight into principles of morphogenesis but potentially also human disease.

## MATERIALS AND METHODS

### Zebrafish

AB, TU and WIK strains of *Danio rerio* were used. Zebrafish lines generated were *sspo*^*b1446*^, *urp1*^*b1420*^ (called *urp1*^*ΔP*^), *urp2*^*b1421*^ (called *urp2*^*ΔP*^), *uts2r3*^*b1436*^ as well as previously published lines: *cfap298*^*tm304*^ (Jaffe et al., 2016), *pkd2l1*^*icm02*^ (Sternberg et al., 2018) and *sspo-gfp*^*ut24*^ (Troutwine et al., 2020).

### Generation and genotyping of mutant lines

*sspo*^*b1446*^, *urp1*^*b1420*^, *urp2*^*b1421*^ and *uts2r3*^*b1436*^ mutant lines were generated using CRISPR/Cas9. gRNA oligos were designed using CRISPRscan (Moreno-Mateos et al., 2015). gRNA templates (IDT) were assembled by annealing and extension with Bottom strand ultramer_1 (Table S2) using Taq Polymerase (NEB, M0273) with cycling parameters of 95^°^C (3 min), 95^°^C (30 s), 45^°^C (30 s), 72^°^C (30 s), 72^°^C (10 min) with 30 cycles of the middle three steps. PCR product was purified (Zymo DNA Clean and Concentrator Kit, D4013) then used for *in vitro* RNA synthesis using a MEGAshortscript T7 Transcription Kit (ThermoFisher, AM1354). Synthesized gRNAs were purified (Zymo RNA Clean and Concentrator Kit, R1013) then 150 pg along with 320 pg/nl Cas9 (IDT, 1081058) were injected into one-cell stage fertilized eggs. The mosaic mutant fish resulting from these injections (F_0_ fish) were raised and outcrossed to AB wild-types and DNA was extracted from the resulting F_1_ embryos. Mutant alleles were screened by PCR coupled with restriction enzyme digestion and/or Sanger sequencing (GeneWiz). Embryos from F_1_ clutches harboring mutations were raised to adulthood and outcrossed to AB wild-types to generate F_2_ families which were screened for mutations and raised. The nature of mutations was identified by sequencing DNA extracted from adult fin clips of F_2_ heterozygous fish using CRISPR-ID to deconvolute (Dehairs et al., 2016) and confirmed by sequencing DNA of F_3_ homozygous embryos. *urp1*^*b1420*^ mutants contain a 279 bp deletion and 1 bp insertion that was genotyped by PCR amplification with *urp1_geno_1* and *urp1_geno_2* primers which generates a 460 bp band from wild-type DNA and a 184 bp band from mutant DNA.*urp2*^*b1421*^ mutants harbor a 61 bp deletion and were genotyped by PCR amplification with *urp2*_*geno_1* and *urp2_geno_2* primers followed by gel electrophoresis to distinguish the 283 bp wild-type band and the 226 bp mutant band. *uts2r3*^*b1436*^ mutants contain a 534 bp deletion and were also genotyped by PCR, using *uts2r3_geno_1* and *uts2r3_geno_2*, in which wild-type sequence led to an 832 bp band and mutant sequence a 298 bp band.

The nature of the *sspo*^*b1446*^ mutation was determined by whole genome sequencing. DNA was extracted from mutant embryos using a phenol/chloroform procedure. Libraries were prepared using the FS DNA Library Prep Kit for Illumina sequencing (NEB, E7805). DNA was digested into 150 bp fragments and paired-end sequencing was performed using a NovaSeq 6000 Sequencing System. Trimmomatic (version 0.36) [ILLUMNIACLIP: TruSeq3-PE-2.fa:2:30:10:1:true LEADING:3 TRAILING:3 SLIDINGWINDOW:5:20 MINLEN:42 AVGQUAL:30] was used to remove Illumina adaptor sequences from paired-end reads. Illumina short-read sequences were then aligned to the GRCz11 reference sequence of chromosome 24 using BWA-MEM (version 0.7.01). SAMtools (version 1.8) was used to sort and index reads. Aligned reads in BAM format were analyzed in IGV (version 2.13.1). Mutants were routinely genotyped by PCR amplification with oligos *sspo_geno_1* and *sspo_geno_2*, followed by BsaI restriction digestion to produce 300 bp and 99 bp bands from wild-type DNA and a protected 399 bp band from mutant DNA.

### Generation of somatic mosaic F_0_ mutants (crispants)

Four gRNA oligos per gene for *cfap298, sspo, urp1*, and *urp2* were chosen from a look-up table (Table S2; Wu et al., 2018). gRNAs were synthesized from oligos in multiplex. After being pooled at (10 µM), oligos were annealed and extended with Bottom strand ultramer_2 using Phusion High-Fidelity PCR Mastermix (NEB, M0531) with Phusion High-Fidelity DNA Polymerase (NEB, M05030) using incubations: 98^°^C (2 min), 50^°^C (10 min), 72^°^C (10 min). Assembled oligos were purified and used as templates for *in vitro* RNA synthesis, as described in “Generation and genotyping of mutant lines” section. For mutagenesis, 1000 pg of gRNAs along with 1600 pg/nl Cas9 (IDT, 1081058) were injected into one-cell-stage embryos.

### Quantitation of body curvature at 1-2 dpf

Zebrafish larvae at 28-30 hpf were imaged using a Leica S9i stereomicroscope with integrated 10-magapixel camera. Body angles were calculated using ImageJ (Schindelin et al., 2012) as described in Bearce et al., 2022.

### Quantitative reverse transcriptase PCR (qRT-PCR)

RNA was extracted using a Zymo Direct-Zol RNA Miniprep kit (Zymo Research, R2051). cDNA was prepared from 25 ng of RNA using oligoDT primers in a 20 µl reaction using a High Capacity cDNA Reverse Transcription Kit (ThermoFisher, 4368814). qRT-PCR reactions were performed in real time using 5 µl PowerUp SYBR Green Master Mix (ThermoFisher, A25741), 0.8 µl of 10 µM forward and reverse primers, 1.4 µl of nuclease-free water and 2 µl of diluted cDNA. PCR was performed using a QuantStudio Real Time PCR System (Applied Biosystems) with cycling parameters: 50°C (2 mins), 95°C (10 mins) then 40 cycles of 95°C (15 s) and 60°C (1 min). Each reaction was performed in quadruplicate. Quantitation was relative to *rpl13* and used the ΔΔC_T_ relative quantitation method in which fold changes are calculated as 2^-ΔΔCT^. The efficiency of amplification was verified to be close to 100% with a standard curve of RNA dilutions.

### Calcein staining

Larvae were incubated in water containing 0.2% calcein (Sigma-Aldrich, C0875) for 10 min then rinsed 2-3 times in water (5 min per rinse). Larvae were immobilized with 0.005% tricaine, mounted in 0.8% low melt agarose and imaged with a Leica THUNDER stereoscope.

### Multiplex fluorescent *in situ* hybridization chain reaction (*in situ* HCR)

Embryos were fixed in 4% paraformaldehyde at 4°C overnight, washed with phosphate buffered saline (PBS) then serially dehydrated to 100% methanol and stored at -20°C. Embryos were rehydrated, washed with PBS, incubated in hybridization buffer (Molecular Instruments), then incubated in 2 pg of probes at 37°C overnight in a total volume of 500 µl of hybridization buffer. Embryos were washed in wash buffer (Molecular Instruments), then incubated in amplification buffer (Molecular Instruments) for 1 hour. RNA hairpins designed to bind either *pkd2l1, urp1* or *urp2* were prepared by heating 10 pmol of each to 95°C for 90 s then snap-cooled in the dark for 30 mins. Embryos were then incubated overnight in 500 µl of amplification buffer containing 10 pmol hairpins at room temperature in the dark. Embryos were washed 5 times in 5 x SSCT, stored at 4^°^C then mounted for confocal microscopy. Images were acquired using a Zeiss LSM880 using either a 20X air or 40X water objective. Acquisition settings were derived using wild type embryos and then applied to all embryos. Images were exported to IMARIS 9.9 (Oxford Instruments). A Gaussian filter of width 0.21 µm (20X) or 0.42 µm (40X) and a rolling ball background subtraction of 10 µm was applied.

### X-ray microcomputed tomography (µCT)

Scans were performed using a vivaCT80 (Scanco Medical) at 18.5 µm voxel resolution as previously described (Bearce et al., 2022).

### Live imaging of SCOspondin-GFP

Embryos (28 hpf) and larvae (12 dpf) were anesthetized in tricaine until touch response was abolished then embedded in 0.8% low-melt agarose laced with tricaine in inverted imaging chambers (14 mm #1.5 coverslips, VWR cat no. 10810-054). In larvae, care was taken to align the posterior body close to the coverslip to the the RF within the working distance of the objective. A Nikon Ti2 inverted microscope equipped with Plan Apo 40X and 60X WI DIC (1.2 NA) objectives, a Yokogawa Spinning Disk and pco.edge sCMOS camera were used to capture 512×256 images in time series. Exposure time varied with age (100 ms - 300 ms) as Sspo-GFP brightened in intensity over time; exposure, camera settings and laser power were kept constant between age-matched individuals. Images were cropped, rotated and intensity-adjusted in ImageJ (Schindelin, 2012).

## ACKNOWLEDGEMENTS

We thank Judy Peirce, Tim Mason and the Aquatics Facility for zebrafish husbandry, the GC3F Biological Imaging Facility and the X-Ray Imaging Core, all at the University of Oregon. We thank Claire Wyart, John Postlethwait, and Ryan Gray for sharing zebrafish lines, Zac Bush for help with genome sequencing, Mike Harms and Ron Kwon for discussions and Katie Fisher for proof reading.

## Author contributions

DTG conceived the project. EAB performed experiments and quantitative analyses. ZHI generated zebrafish mutant lines. JOS performed phenotyping experiments, in situ HCR and quantitative PCR. CJK performed zebrafish injections and imaging. SIF performed zebrafish imaging time courses and Cobb angle measurements. WEC performed quantitative PCR. DTG wrote the manuscript with input from EAB, ZHI and JOS.

