## Supplementary File for "Urotensin II-Related Peptides, Urp1 and Urp2, Control Zebrafish Spine Morphology"

**File inventory:**

Supplementary figures

Fig. S1 | Generation of *urp1*$^{\Delta P}$ and *urp2*$^{\Delta P}$ mutants
Fig. S2 | Generation of *sspo*$^{b1446}$ and *uts2r3*$^{b1436}$ mutants
Fig. S3 | Phenotyping spinal curve mutants

Supplementary tables

Table S1 | Key Resource Table
Table S2 | Oligonucleotides

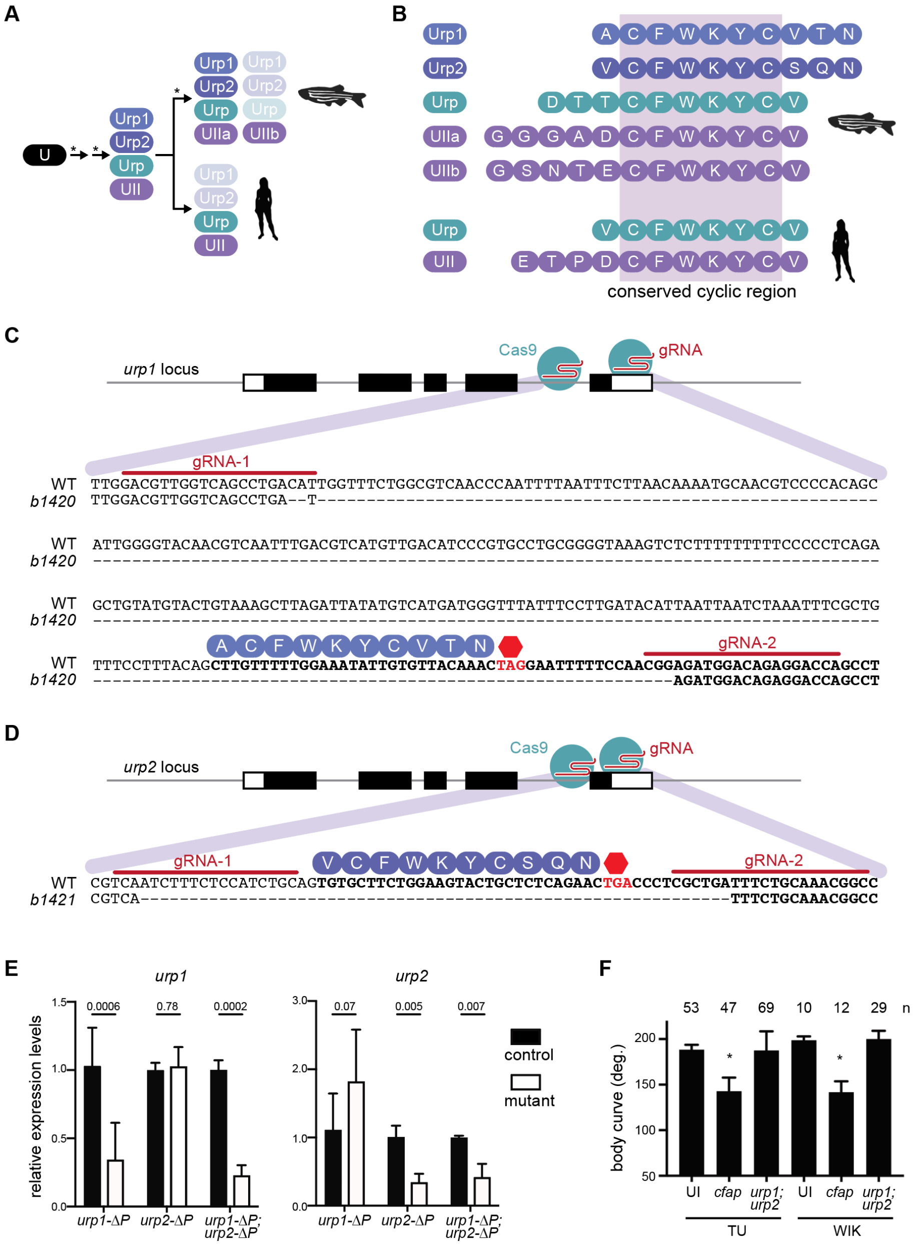


**Fig. S1 | Generation of *urp1^ΔP^* and *urp2^ΔP^* mutants. A)** Successive rounds of genome duplications (*) and divergence converted an ancient Urotensin protein (U) into the Urotensin II (UII) and Urotensin II-related (URP) proteins, some of which have subsequently been lost. **B)** The UII and URP proteins in zebrafish and human are 8-12 amino acid peptides with a fully conserved hexacyclic region of sequence CFWKYC. **C-D)** Pairs of guide RNAs (gRNAs) were used to delete genomic regions coding for the Urp1 and Urp2 peptides. The $urp1^{b1420}$ ($urp1^{\Delta P}$) allele encodes a 279 base pair deletion and 1 base pair insertion that removes a portion of intron 4-5 and the coding part of exon 5 including the entire region coding for the Urp1 peptide. The $urp2^{b1421}$ ($urp2^{\Delta P}$) allele encodes a 61 base pair deletion which removes the region coding for the Urp2 peptide. **E)** Quantitative reverse transcriptase PCR (qRT-PCR) analysis of *urp1* and *urp2* mRNA expression levels in $urp1^{\Delta P}$ and $urp2^{\Delta P}$ single and double mutants at 28 hpf. n >3 biologically independent samples. Bars represent mean ± s.e.m. Twotailed student's *t* test used to calculate *P* values. **F)** Quantitation of crispant body curves where bars represent mean ± s.d. for at least three independent clutches and injection mixes. The total number of embryos analyzed is given. *$P$ < 0.0001, student's *t* test applied. UI – uninjected; *cfap* - *cfap298*.

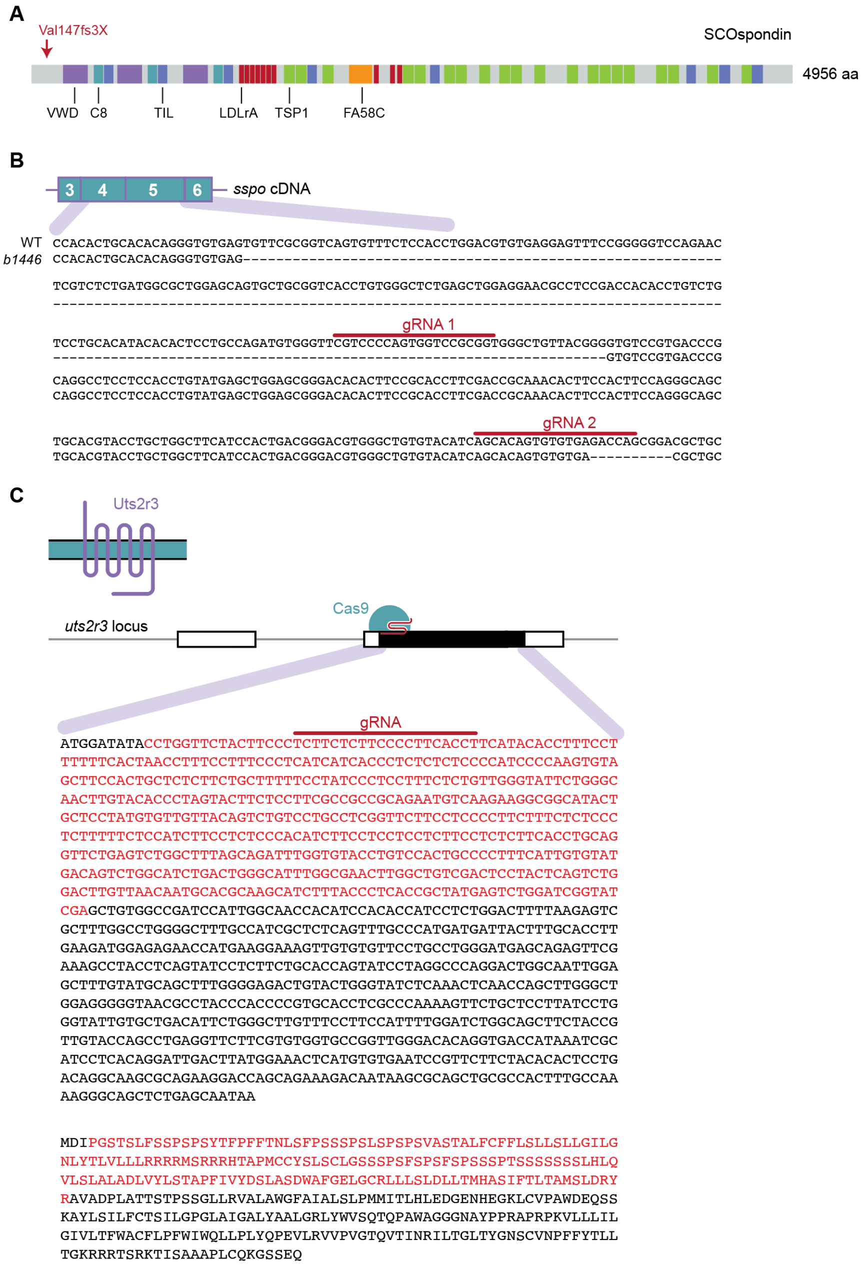


**Fig. S2 | Generation of $sspo^{b1446}$ and $uts2r3^{b1436}$ mutants. A)** Schematic of zebrafish SCOspondin protein showing domain architecture based on Troutwine et al., 2020. VWD – von Willebrand factor type D domain; C8 – C8 domain; TIL – Trypsin Inhibitor like cysteine rich domain; LDLrA – Low Density Lipoprotein Receptor Class A domain; TSP1 – Thrombospondin type 1 domain; FA58C – Coagulation factor 5/8 C-terminal domain. In $sspo^{b1446}$ mutants, a genomic deletion results in a frame shift mutation at Valine 147 resulting in an early premature truncation codon. **B)** The $sspo^{b1446}$ mutant line harbors a large deletion and a downstream small deletion which disrupts exons 4 and 5 causing the early truncation of Sspo. **C)** The $uts2r3^{b1436}$ allele was generated with a single guide RNA which induced a deletion of 534 base pairs, resulting in an in-frame 178 amino acid deletion that removes around half the protein including transmembrane regions.

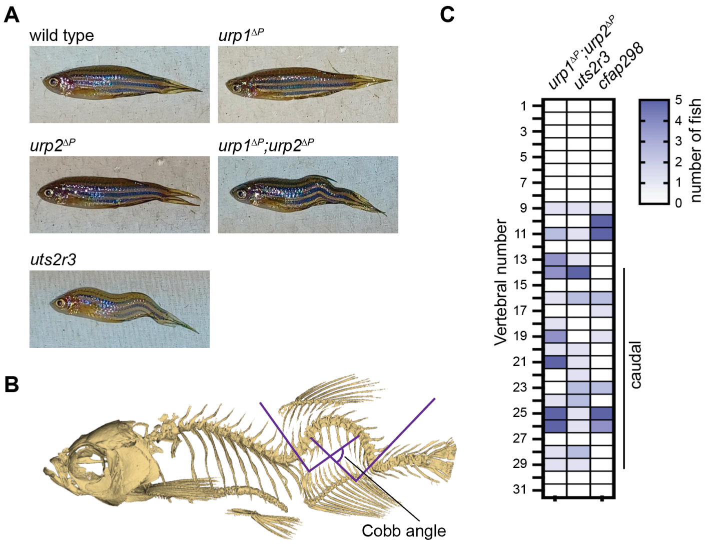


**Figure S3. Phenotyping spinal curves.** **A)** Lateral views of adult zebrafish. **B)** Cobb angles were measured from lateral views of μCT reconstitutions by drawing two lines parallel to the most displaced vertebrae either side of the curve. The Cobb angle was then taken to be the angle made between lines perpendicular to those two lines. **C)** The position of curve apex is plotted for *cfap298$^{tm304}$* mutants alongside *urp1$^{\Delta P}$;urp2$^{\Delta P}$* and *uts2r3$^{b1436}$* mutants for comparison. See also Fig. 2H.

### Table S1 | Key Resource Table

| REAGENT or RESOURCE | SOURCE | IDENTIFIER |
|---|---|---|
| **Commercial Assays** | | |
| DNA Clean and Concentrator Kit | Zymo Research | Cat no: D4013 |
| RNA Clean and Concentrator Kit | Zymo Research | Cat no: R1016 |
| Direct-zol RNA MiniPrep Kit | Zymo Research | Cat no: R2050 |
| GeneJET Gel Extraction Kit | Thermo Fisher Scientific | Cat no: K0691 |
| High Capacity RNA-to-cDNA Kit | Thermo Fisher Scientific | Cat no: 4387406 |
| MEGAshortscript T7 Transcription Kit | Thermo Fisher Scientific | Cat no: AM1354 |
| HiScribe T7 High Yield RNA Synthesis Kit | New England Biolabs | Cat no: E2040 |
| FS DNA Library Prep Kit for Illumina | New England Biolabs | Cat no: E7805 |
| HCR-RNA FISH Hybridization Buffer | Molecular Instruments | |
| HCR-RNA FISH Amplification Buffer | Molecular Instruments | |
| AlexaFluor-647 Hairpins | Molecular Instruments | |
| AlexaFluor-546 Hairpins | Molecular Instruments | |
| AlexaFluor-488 Hairpins | Molecular Instruments | |
| *urp1* HCR-RNA FISH probe | Molecular Instruments | Project-specific design |
| *urp2* HCR-RNA FISH probe | Molecular Instruments | Project-specific design |
| *pkd2l1* HCR-RNA FISH probe | Molecular Instruments | Project-specific design |
| **Chemicals, Peptides and Recombinant Proteins** | | |
| Taq Polymerase | New England Biolabs | Cat no: M0273 |
| TURBO DNase | Thermo Fisher Scientific | Cat no: AM2238 |
| Phusion High-Fidelity DNA Polymerase | New England Biolabs | Cat no: M0530 |
| Phusion High-Fidelity PCR Master Mix | New England Biolabs | Cat no: M0531 |
| SYBR Green PCR Master Mix | Thermo Fisher Scientific | Cat no: 4309155 |
| **Experimental Models: Organisms** | | |
| Zebrafish (*Danio rerio*), AB strain | University of Oregon | |
| Zebrafish (*Danio rerio*), WIK strain | University of Oregon | |

| REAGENT or RESOURCE | SOURCE | IDENTIFIER |
|---|---|---|
| Zebrafish (*Danio rerio*), TU strain | University of Oregon | |
| *cfap298*$^{tm304}$ line | Jaffe et al., 2016 | ZDB-FISH-150901-23024 |
| *pkd2l1*$^{icm02}$ line | Sternberg et al., 2018 | ZDB-FISH-160811-9 |
| *sspo*$^{b1446}$ line | This study | |
| *sspo-GFP*$^{ut24}$ line | Troutwine et al., 2020 | ZDB-FISH-190313-20 |
| *urp1*$^{b1420}$ line | This study | |
| *urp2*$^{b1420}$ line | This study | |
| *uts2r3*$^{b1436}$ line | This study | |
| **Software** | | |
| 3D Slicer | Fedorov et al., 2012 | |
| IMARIS 9.9 | Oxford Instruments | |
| NIS-Elements | Nikon Instruments Inc | |
| ZEN Software | Carl Zeiss AG | |
| QuantStudio Design and Analysis Software | Applied Biosystems | |

### Table S2 | Oligonucleotides

All oligonucleotides were purchased from Integrated DNA Technologies, Inc., USA

| **Name** | **Sequence (5'-3')** | **Function** |
|---|---|---|
| *pkd2l1_geno_1* | TGTGTGCTAGGACTGTGGGG | *pkd2l1*$^{icm02}$ genotyping oligo 1 |
| *pkd2l1_geno_2* | AGGGCAAGAGAATGGCAAGACG | *pkd2l1*$^{icm02}$ genotyping oligo 2 |
| *urp1_gRNA_1* | taatacgactcactataGGCGTTGGTCAGCCTGACATgttttagagctagaa | gRNA_1 oligo for generating *urp1*$^{b1420}$ line |
| *urp1_gRNA_2* | taatacgactcactataGGGTCCTCTGTCCATCTCCGgttttagagctagaa | gRNA_2 oligo for generating *urp1*$^{b1420}$ line |
| *urp1_geno_1* | GCACCCAAAATCCAACGACT | *urp1*$^{b1420}$ genotyping oligo 1 |
| *urp1_geno_2* | TGTATGGGGAAAACAAAGGCA | *urp1*$^{b1420}$ genotyping oligo 2 |
| *urp2_gRNA_1* | taatacgactcactataGGCAGATGGAGAAAGATTGAgttttagagctagaa | gRNA_1 oligo for generating *urp2*$^{b1421}$ line |
| *urp2_gRNA_2* | taatacgactcactataGGCGTTTGCAGAAATCAGCGgttttagagctagaa | gRNA_2 oligo for generating *urp2*$^{b1421}$ line |
| *urp2_geno_1* | TTGGGGTTGTAACAGGTAGTG | *urp2*$^{b1421}$ genotyping oligo 1 |
| *urp2_geno_2* | AACAAGGAAGACGCTGCAAG | *urp2*$^{b1421}$ genotyping oligo 2 |
| *uts2r3_gRNA* | taatacgactcactataGGGTGAAGGGGAAGAGAAGAgttttagagctagaa | gRNA oligo for generating *uts2r3*$^{b1436}$ line |
| *uts2r3_geno_1* | ATGGATCCCCTGATGTCCTG | *uts2r3*$^{b1436}$ genotyping oligo 1 |
| *uts2r3_geno_2* | TCGAACTCTGCTCATCCCAG | *uts2r3*$^{b1436}$ genotyping oligo 2 |
| *sspo_gRNA_1* | taatacgactcactataGGTCCCCAGTGGTCCGCGGTgttttagagctagaa | gRNA_1 oligo for generating *sspo*$^{b1446}$ line |
| *sspo_gRNA_2* | taatacgactcactataGGCACAGTGTGTGAGACCAGgttttagagctagaa | gRNA_2 oligo for generating *sspo*$^{b1446}$ line |
| *sspo_geno_1* | CGCAAACACTTCCACTTCCA | *sspo*$^{b1446}$ genotyping oligo 1 |
| *sspo_geno_2* | TTGAAGCCAGATGTAAAGGATGAGTGT | *sspo*$^{b1446}$ genotyping oligo 2 |
| *urp1_F0_gRNA_1* | TAATACGACTCACTATAGGAAAGTGAAGATCGCGGCCGTTTTAGAGCTAGAAATAGC | gRNA_1 oligo for generating *urp1* F0 embryos |
| *urp1_F0_gRNA_2* | TAATACGACTCACTATAGGACACGGCTCTGCCACAACGTTTTAGAGCTAGAAATAGC | gRNA_2 oligo for generating *urp1* F0 embryos |
| *urp1_F0_gRNA_3* | TAATACGACTCACTATAGGTTCAGAAGCTGGTAGCAGGTTTTAGAGCTAGAAATAGC | gRNA_3 oligo for generating *urp1* F0 embryos |
| *urp1_F0_gRNA_4* | TAATACGACTCACTATAGGGAAAATAAATAACATGGTGTTTTAGAGCTAGAAATAGC | gRNA_4 oligo for generating *urp1* F0 embryos |

| | | |
|---|---|---|
| *urp2_F0_gRNA_1* | TAATACGACTCACTATAGGTGACTGTCGCTTCAATCGGTTTTAGAGCTAGAAATAGC | gRNA_1 oligo for generating *urp2* F0 embryos |
| *urp2_F0_gRNA_2* | TAATACGACTCACTATAGGGACATTTCCTGACGGAGAGTTTTAGAGCTAGAAATAGC | gRNA_2 oligo for generating *urp2* F0 embryos |
| *urp2_F0_gRNA_3* | TAATACGACTCACTATAGGTGGACACGAGGAGACCGAGTTTTAGAGCTAGAAATAGC | gRNA_3 oligo for generating *urp2* F0 embryos |
| *urp2_F0_gRNA_4* | TAATACGACTCACTATAGGTCACCAGGTAGTGACGGAGTTTTAGAGCTAGAAATAGC | gRNA_4 oligo for generating *urp2* F0 embryos |
| *sspo_F0_gRNA_1* | TAATACGACTCACTATAGGTTCGTCCCCAGTGGTCCGGTTTTAGAGCTAGAAATAGC | gRNA_1 oligo for generating *sspo* F0 embryos |
| *sspo_F0_gRNA_2* | TAATACGACTCACTATAGGAAACGGCCGTCAGTGTCGGTTTTAGAGCTAGAAATAGC | gRNA_2 oligo for generating *sspo* F0 embryos |
| *sspo_F0_gRNA_3* | TAATACGACTCACTATAGGTGTTGCAACACCAACCGGGTTTTAGAGCTAGAAATAGC | gRNA_3 oligo for generating *sspo* F0 embryos |
| *sspo_F0_gRNA_4* | TAATACGACTCACTATAGGAGCCTAGACCTGCTCACGGTTTTAGAGCTAGAAATAGC | gRNA_4 oligo for generating *sspo* F0 embryos |
| *cfap298_F0_gRNA_1* | TAATACGACTCACTATAGGTTCTCTTCAACACTACGGGTTTTAGAGCTAGAAATAGC | gRNA_1 oligo for generating *cfap298* F0 embryos |
| *cfap298_F0_gRNA_2* | TAATACGACTCACTATAGGGCTCCACAATCTGATCATGTTTTAGAGCTAGAAATAGC | gRNA_2 oligo for generating *cfap298* F0 embryos |
| *cfap298_F0_gRNA_3* | TAATACGACTCACTATAGGCATTCTTATTGGATCATGGTTTTAGAGCTAGAAATAGC | gRNA_3 oligo for generating *cfap298* F0 embryos |
| *cfap298_F0_gRNA_4* | TAATACGACTCACTATAGGTCTCTGGCAGGTGCGCCCGTTTTAGAGCTAGAAATAGC | gRNA_4 oligo for generating *cfap298* F0 embryos |
| *Bottom strand ultramer_1* | AAAAGCACCGACTCGGTGCCACTTTTTCAAGTTGATAACGGACTAGCCTTATTTTAACTTGCTAT | tail ultramer for generating singe gRNA oligos |
| *Bottom strand ultramer_2* | AAAAGCACCGACTCGGTGCCACTTTTTCAAGTTGATAACGGACTAGCCTTATTTTAACTTGCTATTTCTAGCTCTAAAAC | tail ultramer for generating multiplexed gRNA oligos |
